# On the unfounded enthusiasm for soft selective sweeps II: examining recent evidence from humans, flies, and viruses

**DOI:** 10.1101/443051

**Authors:** Rebecca B. Harris, Andrew Sackman, Jeffrey D. Jensen

## Abstract

Since the initial description of the genomic patterns expected under models of positive selection acting on standing genetic variation and on multiple beneficial mutations—so-called soft selective sweeps—researchers have sought to identify these patterns in natural population data. Indeed, over the past two years, large-scale data analyses have argued that soft sweeps are pervasive across organisms of very different effective population size and mutation rate—humans, Drosophila, and HIV. Yet, others have evaluated the relevance of these models to natural populations, as well as the identifiability of the models relative to other known population-level processes, arguing that soft sweeps are likely to be rare. Here, we look to reconcile these opposing results by carefully evaluating three recent studies and their underlying methodologies. Using population genetic theory, as well as extensive simulation, we find that all three examples are prone to extremely high false-positive rates, incorrectly identifying soft sweeps under both hard sweep and neutral models. Furthermore, we demonstrate that well-fit demographic histories combined with rare hard sweeps serve as the more parsimonious explanation. These findings represent a necessary response to the growing tendency of invoking parameter-heavy, assumption-laden models of pervasive positive selection, and neglecting best practices regarding the construction of proper demographic null models.

## INTRODUCTION

For decades, identifying beneficial mutations based on genomic patterns of linked polymorphism has remained a topic of keen interest theoretically, methodologically, and empirically in the field of population genetics. Initial efforts were largely focused around a hard selective sweep model—that is, one in which positive selection acts upon a newly arising beneficial mutation and brings it to fixation in the population (see Maynard Smith & Haigh [1]). Under this model, multiple reasonably well-performing methods have been developed for detecting hard sweeps, though they all commonly struggle with low power and/or high false-positive rates under certain neutral non-equilibrium models (*e.g*., severe population bottlenecks; see review of [2]).

Owing both to theoretical developments (*e.g*., [3,4]), as well as a lack of evidence for widespread hard sweeps in the genomes of commonly studied organisms (*e.g*., [5]), alternative models have gained attention over the past decade. For example, the notion of soft selective sweeps encompasses at least two very different models: a) selection on standing variation—in which positive selection begins acting upon a mutation only once it is already at appreciable frequency in the population, and b) multiple *de novo* beneficial mutations—in which positive selection acts upon independently-arising and simultaneously-segregating copies of (a) beneficial mutation(s). Despite being relevant in two very different parameter spaces, the commonality between these two models and the reason for their common grouping as soft sweeps is that both models may result in multiple high frequency haplotypes at the time of fixation. This reflects the fact that equivalent copies of beneficial mutations are carried on different haplotypes at the onset of selection, and therefore these haplotype backgrounds may hitchhike to intermediate frequencies.

A number of objections have been raised, not against the models themselves *per se*, but against their relative applicability to empirical population data. With regards to the above prediction of multiple high frequency haplotypes, Schrider *et al.* [6] noted that the presence of high frequency haplotypes is also a widely-utilized prediction of a hard sweep model with recombination. Namely, because mutations may recombine on/off the selected haplotype during the course of a selective sweep, and do so with increasing recombination distance, mutations that are partially linked to the beneficial mutation will only be brought to intermediate frequency. As such, the authors described a so-called ‘soft-shoulder effect’ in which regions flanking a hard sweep may be mis-characterized as being the target of a soft sweep.

With regards to selection on standing variation, Orr and Betancourt [7] demonstrated that the likelihood of a hard sweep reaching fixation is not simply determined by whether positive selection begins acting when the beneficial mutation is present in a single vs. multiple copies in the population. Instead, they showed that there is a wide parameter space under which selection on standing variation will still result in a hard sweep (that is, a single haplotype fixed at the target of selection). This likelihood is dependent upon the strength of selection acting on the mutation prior to becoming beneficial (*i.e*., whether it is neutral or weakly deleterious), the strength of selection acting on the mutation after becoming beneficial, and the population mutation rate (though see the argument of Hermisson and Pennings [8] on deterministic vs. stochastic sweep approximations). Regardless, the argument that a mutation which significantly impacts the phenotype (to such an extent that it may be strongly beneficial) is segregating as a relatively high frequency neutral, nearly-neutral, or deleterious mutation prior to the shift in selective pressure remains largely lacking in terms of empirical examples.

With regards to the model of multiple competing beneficial mutations, Jensen [9] examined the conditions under which a soft rather than a hard sweep may result. In this case, the likelihood of a soft sweep inherently relies on: 1) a high beneficial mutation rate or large population size (with some arguing that the relevant population size consideration in this case may be much larger than the effective population size; Garud *et al.* [10]), as a second identical *de novo* beneficial mutation must arise during the sojourn time of the first mutation; 2) a large mutational target size, as the competing *de novo* beneficial mutations must have identical selective effects (lest one out-compete the other and result in fixation of the most beneficial (*i.e.* a hard sweep)); and 3) all sites must be essentially independent of one another (*i.e.,* freely recombining). With regards to the latter, if we condition on the unlikely event that two identical beneficial mutations appear in the population in quick succession, if they sit on different linked haplotypes (as is required for a soft sweep), those haplotype backgrounds are highly unlikely to also be of identical selective effect size. Thus, these conditions are likely nearest to being met in certain high mutation/recombination rate viruses.

Despite these difficulties, the enthusiasm for invoking soft sweeps to explain observed patterns of genomic variation has not waned. Garud *et al*. [10] evaluated populations of *Drosophila melanogaster*, proposing a statistic based on a comparison of haplotype frequencies (termed H12, and H2/H1). With this, they argued for genome-wide selective sweep patterns, characterizing all of the top 50 putatively swept regions as likely to be soft rather than hard sweeps. Similarly, Schrider and Kern [11] argued that soft sweeps are the dominant mode of adaptation in humans. Looking across six human populations, they identified 1,927 distinct selective sweeps patterns, 1,776 of which were classified as being a soft sweep using S/HIC, a supervised machine learning approach developed by the same authors [12]. Finally, evaluating 6,717 HIV-1 consensus sequences from patients sampled between 1989-2013, Feder *et al.* [13] demonstrated that viral population diversity is strongly reduced in patients on effective drug treatments, whereas population diversity was not strongly reduced in the viral populations of patients on less effective treatments. In order to justify that this is not simply a description of the strong viral population bottleneck associated with effective (relative to ineffective) drug treatment, the authors further argued that this pattern is correlated with the number of putative drug resistance mutations. Verbally making the case that there is no evidence of a correlation between the strength of selection acting on these drug resistance mutations and the actual degree of drug resistance conferred, the authors instead argue that ineffective drug treatments are associated with soft sweeps, whereas effective drug treatments are associated with hard sweeps.

Given this confusing literature, we closely investigated the basis of these three recent claims and carefully evaluated the newly proposed methodologies underlying them. In particular, we examined the impact of the population’s demographic history on the inferred mode of selection. We demonstrate that, owing to the wide-range of expected patterns of variation produced under models of soft sweeps, these new methods frequently classify both neutral demographic histories, as well as hard sweeps, as soft sweeps. In short, soft selective sweeps appear as the default conclusion under any model that results in intermediate frequency haplotypes. As such, recent claims for widespread soft sweeps are highly tenuous, and simulation results demonstrate that well-fit demographic models combined with rare hard sweeps stand as a strong alternative explanation for observed data. Given the highly non-equilibrium history characterizing all three examples, a recurrently well-supported distribution of fitness effects characterized by rare beneficial mutations, as well as the model specifics outlined by Jensen [9], we argue that pervasive soft sweeps are a highly unlikely explanation in all instances. As these examples were chosen to span organisms of very different underlying population parameters (*e.g*., effective population size, mutation rate, and demographic history), the results and considerations described below are therefore applicable to the much broader soft selective sweep literature.

## RESULTS & DISCUSSION

We here give treatment in turn to the analyses of Garud *et al*. [10], Schrider and Kern [11], and Feder *et al*. [13], evaluating their underlying methodologies, recapitulating their results, and examining alternative models.

### A detailed look at the methodology and claims of Garud et al. 2015

It is first necessary to examine the performance of the H12 and H2/H1 statistics on which subsequent claims about the mode of selective sweeps in Drosophila are based. Both hard and soft sweeps result in reduced genetic variation and haplotype structure compared to a locus evolving under equilibrium neutrality. Hard sweeps result in the rise of a single high frequency haplotype at the target of selection, whereas soft sweeps result in multiple high frequency haplotypes. To distinguish between neutral and selected regions, Garud *et al*. [10] proposed the H12 statistic, which captures the degree of haplotype homozygosity. By combining the frequencies of all haplotypes into a single statistic, values close to one are indicative of both hard and soft sweeps, whereas smaller values are found with many low frequency haplotypes, which they take as indicative of neutrality. Conditional on the initial ascertainment of regions based on H12, they further parse between hard and soft sweeps from amongst these outliers using an H2/H1 statistic, which represents the frequencies of all but the most common haplotype divided by the frequencies of all haplotypes. The expectation is that soft sweeps will have large values of H2, but small values of H1, leading to a H2/H1 statistic close to one. The reverse is expected to be true for hard sweeps.

While Garud *et al.* [10] demonstrated that the means of these statistics vary across different selection regimes, they neglected to provide the information necessary to determine Type I and Type II error under more realistic demography. Indeed, as outlined in their Methods section, all performance analyses of their haplotype statistics were conducted under equilibrium neutrality. More specifically, they did not explore the ability of the statistics to detect positive selection, or to differentiate hard and soft sweeps, within the context of more realistic non-equilibrium demographic histories. Furthermore, these limited equilibrium power results were presented only for incomplete sweeps (*i.e.,* when the beneficial mutation is at 50% frequency in the population - also shown as 10% and 90% in their Supplementary Materials) - further confounding the ability to interpret hard vs. soft sweep performance. Given the rapid sojourn time of a beneficial mutation, the assumption that all sweeps are on-going is peculiar indeed.

Furthermore, and as discussed in Vy *et al*. [14], *H*-statistics were calculated using a fixed window size. This is problematic given that the size of the region affected by a sweep is proportional to both the strength of selection as well as the recombination rate. Under a scenario of weak positive selection or high recombination rate, the fully swept genomic region may be smaller than the window size. In this example, recombination would be expected to lead to multiple haplotypes within a window centered on a hard sweep, resulting in the incorrect inference of a soft sweep (a notion related to the ’soft-shoulder’ effect described by Schrider *et al.* [6]). Indeed, Vy *et al.* demonstrated that using a flexible window-size approach results in hard-sweep classifications for loci identified as soft sweeps by Garud *et al.*

Finally, it is important to note that the ultimate application of these statistics was the Drosophila Genetic Reference Panel (DGRP) dataset, a population of inbred lines from North Carolina, USA. Given extensive evidence in *Drosophila melanogaster* of a complex demographic history, including admixture as well as extensive population size change (*e.g*., [15]), both the demographic models and parameter space explored by Garud *et al.* [10] are of great importance. Indeed, Duchen *et al.* [15] themselves noted the uncertainty of their North American population size inference and, as is typical of any demographic analysis, provided a 95% credibility interval for each of their demographic parameter estimates (see Table 4 & 5 of [15]). Rather than exploring this relevant parameter space, Garud *et al*. [10] generated *H*-statistics for the mode of each parameter. In so doing, they found that the DGRP dataset has uniformly elevated H12 values genome-wide relative to their simulated data (see Fig 7 of [10])—which consisted of two equilibrium models (*N_e_* = 10^6^ and *N_e_* = 10^8^), two relatively old bottleneck models of varying severity, and the point estimates of Duchen *et al*. [15]. They then proceeded to select the top 50 peaks to explicitly test for congruence with a hard vs. soft sweep model, and ultimately claimed that all of these top candidate regions are the product of soft sweeps. Following this logic, however, leads to the fanciful conclusion that the entire Drosophila genome is shaped by positive selection, given that all of the regions tested by Garud *et al.* appear to be H12 outliers (see Fig 7 of [10]).

In order to address the narrow focus on modal demographic parameter values, we firstly accounted for the statistically-described uncertainty in these estimates by rather sampling directly from the 95% credibility intervals of the posterior density for each demographic parameter (S1 Table). For each randomly drawn set of parameters from these posteriors, we generated neutral, hard, and soft sweep simulations. As their methodology relies on initially detecting H12 outliers, we first used simulated data to test the power of the H12 statistic to distinguish neutrality from positive selection under the DGRP demographic model. Similarly, we used simulated data to test whether H2/H1 can discern between hard and soft sweeps under this model. Second, we explored whether the empirical DGRP H12 peaks are indeed outliers compared to the distribution expected under the inferred neutral demographic history.

In so doing, we demonstrate that the *H*-statistics are inadequate to detect or differentiate sweeps under the DGRP demographic model. Matched for identical demographic parameters, neutral simulations produce H12 values greater than or equal to soft sweeps > 53% of the time. In addition, the H2/H1 statistic utilized to discern the mode of selection has poor discriminatory power: hard sweeps have H2/H1 values greater than or equal to soft sweeps > 64% of the time (and see Fig.S1 for a corresponding Bayes Factor plot).

Turning to the empirical DGRP data, Fig 1 demonstrates that the empirically observed H12 outlier values fall within the tail of the distribution expected under the inferred demographic model (*p* = 0.001 - 0.004; corresponding to 16-59 neutral simulations with more extreme H12 values), and are not more extreme than expected under neutrality. In other words, these observations are fully consistent with scanning whole-genome data and ascertaining the most extreme regions (as indeed was done), and as such there is no need to invoke anything other than population history in order to explain empirical observations. Separately, it is clear that the H2/H1 statistic utilized to discern the mode of selection has poor discriminatory power. The empirical *p*-values demonstrate that the top 50 peaks claimed to be the result of soft sweeps all have H2/H1 values that fall well within the distribution of hard sweeps (*p* = 0.1-0.33). Together, and contrary to the author’s claims, these results demonstrate an inability to differentiate neutrality from positive selection under this demographic model, and, even if that were not the case, further demonstrate an inability to distinguish hard from soft sweeps.

**Fig 1.**
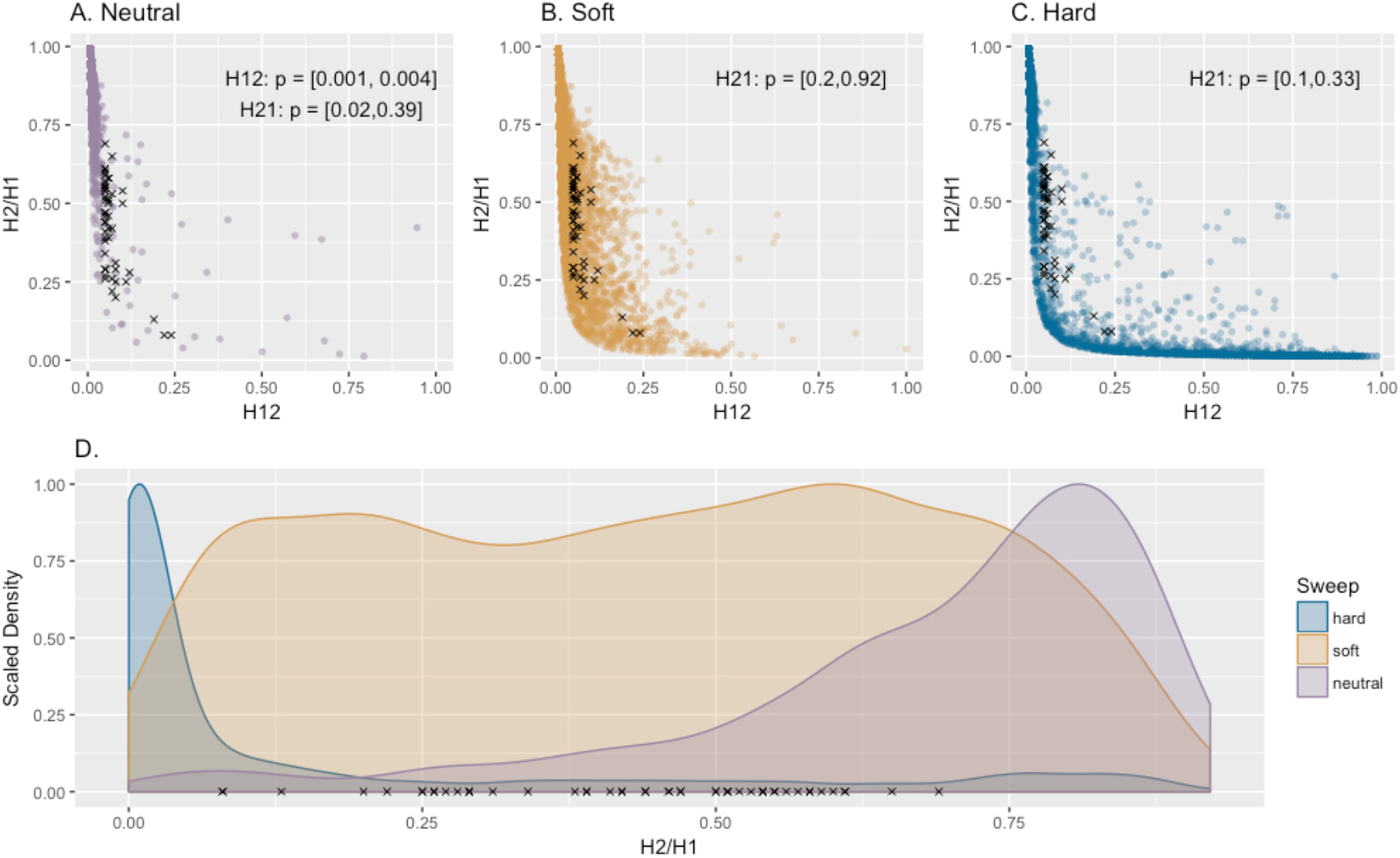
The performance of the *H*-statistics. Distribution of H12 and H2/H1 values estimated under the 95% credibility interval of the DGRP admixture model of Duchen *et al.* [15] for (a) neutrality, (b) soft sweeps, and (c) hard sweeps. Additionally, all panels show the top 50 H12 outliers (black x’s) from the empirical Drosophila data set that Garud *et al.* [10] concluded were soft sweeps. (d) Following their proposed practice, simulations generating the top 2.5% H12 values were ascertained from each set, and the scaled density of the corresponding H2/H1 values are plotted for these H12 outliers. All top 50 empirical outliers fall within the tail of the neutral demographic distribution, as well as within the soft and hard H2/H1 distributions.

In sum, the H12 statistic utilized to ascertain genomic regions becomes elevated under a wide-range of neutral demographic histories, and the H2/H1 statistic utilized to discern the mode of selection from amongst these H12 outliers has poor discriminatory power characterized by largely overlapping distributions between models. Furthermore, and given the above results, when considering a demographic model fit to the DGRP data, it is apparent that the top 50 outlier regions ascertained by Garud *et al.* are fully consistent with neutrality.

### A detailed look at the methodology and claims of Schrider and Kern 2017

It is again necessary to first evaluate the performance of the method itself, S/HIC, a classifier that relies upon a variety of summary statistics (including the above-examined H12 and H2/H1 statistics of Garud *et al.* [10]), before considering the results of their application to human polymorphism data.

S/HIC is a supervised machine learning method which classifies genomic regions into discrete categories (hard, hard-linked, soft, soft-linked, or neutral) by comparing test (empirical or simulated) data to simulated training data. Because demographic perturbations may mask the genomic signals of selection, a user must first choose an appropriate demographic model under which to simulate training data. The initial publication [12] demonstrated that S/HIC has a high true positive rate when the true demographic model is known *a priori*. However, as this is never the case in practice, we examined a range of model mis-specifications. Some of the difficulty in correctly identifying selection under mis-specified demographic models can be found in Fig S10 of Schrider and Kern [12] (and plotted here in Fig 2A). Here, the classifier was trained using an equilibrium demographic model, and the true/test data were simulated under the non-equilibrium model estimated for African human populations by Tennessen *et al.* [16]. Though this Supplementary Figure is cited in the main text as illustrating the ability of S/HIC to robustly deal with unknown non-equilibrium scenarios, when examining the results one finds that even for a hard sweep with extremely strong selection (5000 < 2*Ns* < 50000), the correct model is identified in fewer than 5% of simulated replicates, while hard sweeps are incorrectly classified as soft sweeps in nearly 50% of replicates, as linked soft sweeps in 16% of replicates, and neutrality in 31% of replicates. In other words, under this demographic history estimated for human populations, the original authors demonstrated not only an inability to detect hard sweeps but also an extreme false-positive rate in the direction of mis-classifying them as soft sweeps, if the demographic model is mis-specified.

**Fig 2.**
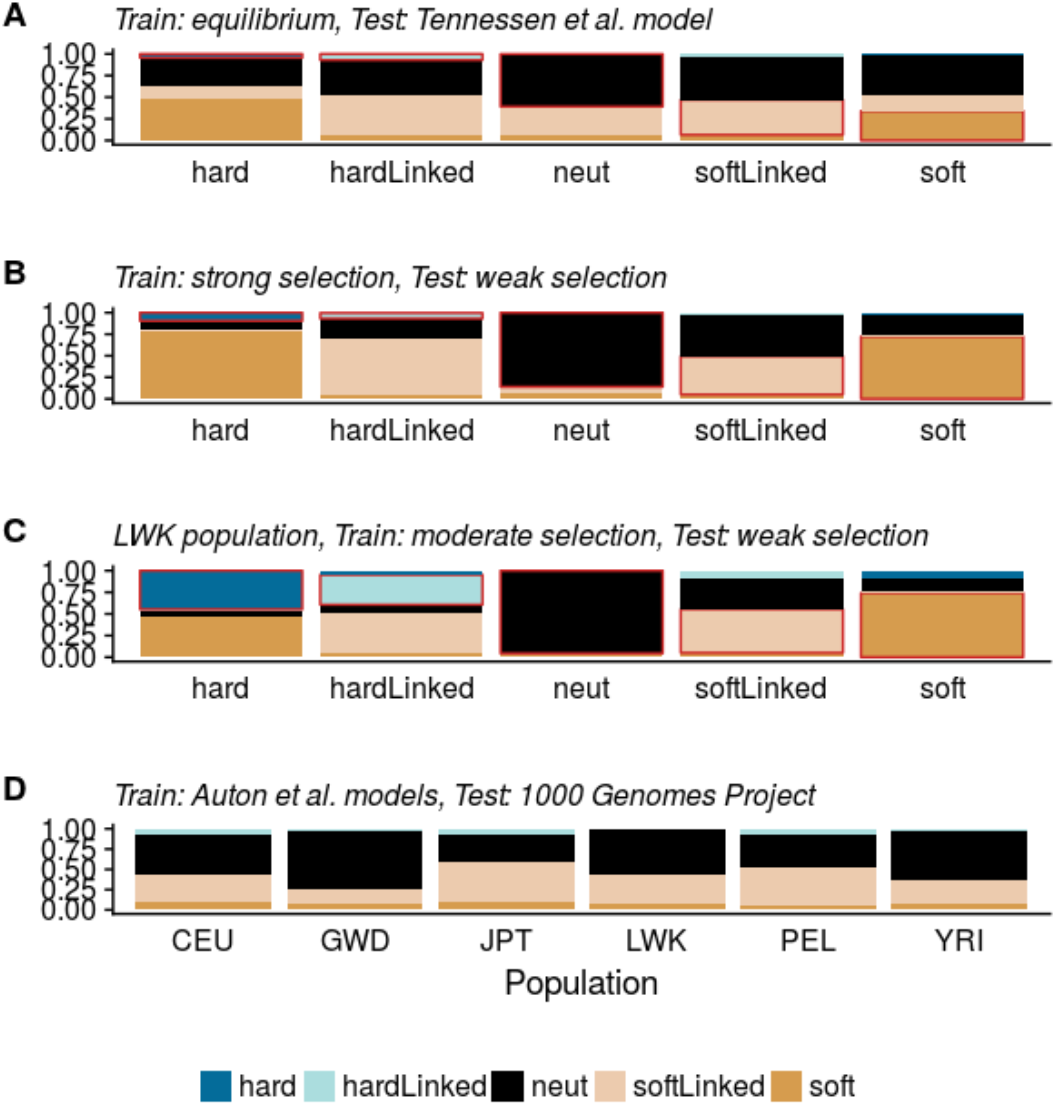
The performance of S/HIC. Stacked bar plots depict the probabilities of model classification by S/HIC. Each vertical bar represents 1000 simulated datasets of each category (where the true models are given on the x-axis in panels a-c (*i.e*., hard, hard-linked, neutral, soft, and soft-linked)). Within each bar, colors represent the proportion that were assigned to each category by S/HIC, and the red outline indicates the correct classifications (*i.e*., true positives). (a) The plotted results of Fig S10 of Schrider and Kern [12], examining classification performance under the Tennessen *et al*. [16] African human demographic model, when the training data assumes an equilibrium model. (b) Results when the strength of selection is mis-specified—both test and training data were simulated under an equilibrium demographic model, where the true dataset is drawn from a moderate selection model (2*Ns* ~ *U*[25, 250]) and the training dataset from a strong selection model (2*Ns* ~ *U*[250, 2500]). (c) Performance when the simulated LWK population experiences weaker selection (2*Ns* ~ *U*[10, 1000]) than the training set (2*Ns* ~ *U*[166, 3333]). (d) The classification ratios of the empirical 1000 Genomes project data presented by Schrider and Kern [11] (also depicted in their Fig 2), which they trained upon a history of population size change as interpreted from PSMC by Auton *et al*. [17]. From left to right on the x-axis, the populations presented are from individuals sampled in North America of Northern and Western European ancestry (CEU), Gambia (GWD), Japan (JPT), Kenya (LWK), Peru (PEL), and Nigeria (YRI).

In presenting results in the main text, the authors depicted the diagnostic ability of binary classifiers (which S/HIC is not), thereby combining classifications (*i.e*., they reported the performance of accurately classifying hard combined with soft sweeps as a single value, and of accurately classifying neutrality combined with linked selection as a single value). Thus, their ROC plots (*e.g*., Fig S9 of [12]) give the appearance that S/HIC performs reasonably well, as they grouped any identification of a soft or hard sweep as a true positive (*i.e*., this grouping neglects the fact that these ’true-positives’ consist of hard sweeps that have been incorrectly identified as soft; or, said another way, that their true positives consist nearly entirely of false positives). Therefore, we differentiate these results in Fig 2, clearly demonstrating widespread mis-classification when the true model is unknown; moreover, this mis-classification is almost universally in the direction of falsely identifying soft selective sweeps.

Even when relatively high probabilities of correct classification are obtained (> 90%), it is critical to examine false positive rates. We note that for a genome the size of humans (~3 billion bp) and scanned using 100 kbp windows (as was done), one would expect thousands of genomic regions to be falsely classified. Therefore, it is of critical importance to compare expected mis-classification rates to observed proportions. In Schrider and Kern [11], S/HIC was used to classify the 1000 Genomes data using the inferred demographic models of Auton *et al*. [17]. Given the null hypothesis that the genome is evolving neutrally, one may multiply the neutral mis-classification rates (reported in their Fig.S1) by the total number of windows analyzed (13,968) to determine how often one should expect a truly neutral region to be mis-classified. In Table 1, we demonstrate that the majority of soft sweeps predicted by Schrider and Kern [11] can be explained as mis-classified neutral regions, given their reported false positive rates.

**Table 1.** The expected number of neutral regions mis-classified as sweeps (soft or hard) compared to the number of sweeps reported by Schrider & Kern [11]. Using the mis-classification proportions reported in their Table S1 and the total number of genomic windows analyzed (*n* = 13,968), one may calculate the number of mis-classified windows expected under neutrality. This represents something of a best case scenario in which the true demographic model is assumed to be known (*i.e.,* that inferred by Auton *et al.* [17]). Comparing these results with those reported by Schrider and Kern [11], it is clear that nearly all of the observed sweep regions can be accounted for by the number of expected false-positives. Populations presented are from individuals sampled in North America of Northern and Western European ancestry (CEU), Gambia (GWD), Japan (JPT), Kenya (LWK), Peru (PEL), and Nigeria (YRI).

| Population | Proportion of neutral regions expected to be mis-classified as sweeps |  | Number of neutral regions expected to be mis-classified as sweeps genome-wide |  | Number of genome-wide sweeps reported by Schrider & Kern 2017 |  |
| --- | --- | --- | --- | --- | --- | --- |
|  | Soft | Hard | Soft | Hard | Soft | Hard |
| CEU | 0.066 | 0.001 | 922 | 14 | 947 | 66 |
| GWD | 0.041 | 0 | 573 | 0 | 795 | 5 |
| JPT | 0.058 | 0.002 | 810 | 28 | 998 | 61 |
| YRI | 0.044 | 0 | 615 | 0 | 797 | 13 |
| PEL | 0.062 | 0.002 | 866 | 28 | 655 | 32 |
| LWK | 0.045 | 0.001 | 629 | 14 | 805 | 3 |

It is notable that additional information is available to potentially improve these classification predictions. The S/HIC pipeline includes a step to estimate the probability that a genomic region belongs to each of the five classes (*i.e*., hard, hard-linked, soft, soft-linked, and neutral), but then only uses this information to rank class labels. As a result, the class label with the highest probability is chosen as the correct model. However, these class probabilities confer critical information pertaining to statistical confidence [18]. In other words, in a scenario where neutral, soft, soft-linked, hard, and hard-linked are classified with 0.19, 0.22, 0.20, 0.19, and 0.19 probabilities, respectively, S/HIC would conclude that the region in question experienced a soft sweep, despite having little support for this classification in reality. Indeed, the number of sweep predictions plummets with the imposition of a probability threshold (Fig.S2). Across all six populations evaluated by Schrider and Kern [11], an average of ~18 soft sweeps were retained with a 0.90 probability threshold. Thus, the inclusion of this uncertainty has the potential to drastically reduced false positive rates.

Similar to the demographic models described above, if the distribution of fitness effects could be known *a priori* (that is, the biologically true underlying selection coefficients are accurately simulated for the training set), S/HIC performs well in classifying models (see Fig 4 of [12]). However, when the training set is simulated with stronger selection than the (true) test set, S/HIC overwhelmingly classifies hard sweep windows as soft (Fig 2B). This result owes to the larger number of recombination events (producing greater haplotypic diversity) and the reduced size of the sweep region in the test set (*i.e*., intermediate strengths of selection) relative to the training set (*i.e*., strong selection). This finding is notable given Schrider and Kern’s assumption that selection coefficients are bounded between 0.005 and 0.1 when analyzing human data [11]. Indeed, when testing a weaker range of selection (2*Ns* ~ *U*[10, 1000]) against that used by Schrider and Kern (2*Ns* ~ *U*[166, 3333]), 47% of hard sweeps are mis-classified as soft, and 46% of hard-linked sweeps are mis-classified as soft-linked (Fig 2C).

Finally, we explored a range of demographic model mis-specifications in order to quantify downstream effects on selection inference. Beginning with population bottlenecks, we considered a scenario in which the applicability of a bottleneck model is known, but the severity of that bottleneck is mis-inferred. If the training set is simulated from a single incorrect bottleneck value, neutral simulations are increasingly inferred to be soft as the true bottleneck size becomes more severe (Fig.S3). However, as demonstrated in the previous section, best-practice would necessitate sampling from across the parameter’s posterior distribution. In order to mimic such a scenario, we constructed training sets of varying widths, and found that this may indeed ameliorate the situation to some extent, though false positive rates in the direction of soft sweeps remain unacceptably high (Fig.S3). In addition, severe population bottlenecks, even when accurately known *a priori*, remain extremely difficult to distinguish from selection.

Next considering hidden population structure, the signatures of genome-wide haplotype distributions associated with such sampling resulted primarily in the detection of soft sweeps regardless of the underlying model (Fig.S3), a pattern that is only exacerbated under models of rare migration (shown for a divergence time of 4*N* generations; Fig.S3 and S2 Table). Given that both structure and migration are highly relevant in the demographic history of humans, we further considered two divergent populations experiencing recent, punctuated gene flow - varying the depth of divergence, the rate of migration, and the number of admixture events. As shown in Fig.S3 and S2 Table, these population dynamics also lead to widespread mis-classification of neutral loci as soft sweeps. These findings are particularly relevant as recent studies have demonstrated that PSMC-based human demographic inference— which neglects a consideration of population structure—generates artificial signals of population size change [19]. Thus, the PSMC patterns interpreted as fluctuations in population size can be replicated by simulations of structured populations experiencing shifts in rates of migration [19,20].

In sum, as with the Drosophila analysis, the claim of soft sweeps in human populations is based on the observation of an excess of intermediate frequency haplotypes across the genome relative to neutral equilibrium expectations. The accurate performance of the statistic relies on a prior knowledge of both the distribution of fitness effects as well as the demographic history of the population in question. As neither is ever accurately known in practice, it is troubling that the mis-specification of either results in pervasive mis-classification. Namely, hard sweeps of weakly beneficial mutations will be classified as soft sweeps; and neutral demographic models which result in haplotype structures similar to soft sweeps (including mild bottlenecks, structured populations, and migration) will be classified as ’soft’, while severe population bottlenecks may result in genomic patterns of variation which appear ’hard’. Furthermore, if the true demographic model is unknown, and considering the Tennessen *et al*. [16] estimates as an example, one finds no power to detect hard sweeps, and an extreme false-positive rate in the direction of mis-classifying them as soft sweeps.

Thus, the repeated claim that S/HIC is robust to demography [11,12,21] is unwarranted. Further, the finding of genome-wide soft sweeps appears entirely consistent with mis-inference owing to both the highly non-equilibrium history of these human populations as well as the underlying assumption of large selection coefficients. Indeed, Schrider and Kern’s average inferred fractions for the genome-wide empirical human data of 0.004 hard sweep windows, 0.045 linked hard sweep windows, 0.084 soft sweep windows, 0.516 linked soft sweep windows, and 0.352 neutral windows matches well the expected mis-classification under a variety of the models illustrated in Fig 2 and S3 Fig. In addition, simply considering reported false positive rates under neutrality, along with the number of genomic windows evaluated across the human genome, is largely sufficient to replicate their reported results.

### A detailed look at the methodology and claims of Feder et al. 2016

In contrast to the above studies, Feder *et al*. [13] do not propose a novel method, but instead rely on the expectation that hard sweeps will reduce variation much more strongly than soft sweeps. It is thus necessary here to examine the specific support for soft and hard sweep models in these HIV-1 patient samples. The authors utilized an analysis of sequence variability to compare the change in diversity following between zero and four sweeps of drug-resistant mutations (DRMs) in a variety of HIV drug treatments of varying degrees of effectiveness. Specifically, they analyzed the consensus HIV-1 population sequence data from the reverse-transcriptase and protease genes for 6,717 patients sequenced over a 24-year span and treated with exactly one drug regimen, with sampling being taken after initiation of treatment (but not necessarily after treatment failure). In Feder *et al*. [13], ambiguous base calls present in the consensus sequence were summed across each sequence and used as a proxy measure of genetic diversity within the population. The authors demonstrated that on average the number of ambiguous base calls per sequence decreased with the addition of each of the first four DRMs, as matched against the 2009 World Health Organization list of DRMs used for surveillance of transmitted HIV-1 drug resistance [22]. Additionally, they successfully demonstrated that the change in diversity accompanying each DRM present in a sequence was significantly correlated with treatment effectiveness, measured as the percentage of patients exhibiting virologic failure after 48 weeks of treatment. The authors relied on this result to argue that less effective treatments exhibit patterns consistent with soft sweeps, whereas more effective treatments exhibit decreases in diversity consistent with hard sweeps. We evaluate here whether soft sweep models actually need be invoked to explain this data.

We implemented a model wherein a sequence of length 1980 nt (equivalent to the combined lengths of the reverse-transcriptase and protease genes) was allowed to evolve in a manner approximating a period of initial infection and exponential population growth, followed by a treatment-induced bottleneck, sequential fixation of beneficial DRMs, and gradual recovery to the pre-treatment population size (see Methods). First, we considered the neutral demographic history characterizing these viral population samples, given the differences in effectiveness of the administered drug treatments across temporally-sampled patients. Though not considered/modeled in Feder *et al*. [13], it is evident that effective treatment strategies translate to a strong reduction in viral population sizes, whereas ineffective treatments do not. Variation in the severity of bottlenecks during treatment must therefore result in differing levels of neutral genetic variability within populations exposed to treatments of differing efficacy. Additionally, populations exposed to more effective treatments that exhibit longer periods of virologic suppression will on average spend longer periods of time at reduced population size.

We first applied our model of HIV infection in 1,000 replicate simulations of populations experiencing sequential fixation of DRMs of identical selection coefficients under three different demographic scenarios: bottlenecks to 10, 10^2^, or 10^3^ genomes at the time of treatment, from an original population size of 10^4^ genomes. These scenarios represent treatments of high, intermediate, and low efficacy, resulting in a range of bottlenecks from more to less severe, respectively. As expected, there is a large and highly significant (*p* << 0.001) difference in the overall level of diversity, measured as the number of ambiguous base calls, between populations experiencing these differing bottleneck severities (Fig 3A). Namely, each ten-fold decrease in population size during the treatment period resulted in at least a ten-fold decrease in the remaining variation after resistance was acquired and populations rebounded. Thus, the assumption of Feder *et al*. [13] that treatments of variable efficacy do not induce population bottlenecks of varying severity could lead to the erroneous conclusion that resistance to weaker treatments must have evolved via soft sweeps, as this is the only model that they considered which would allow for this preservation of variation.

**Fig 3.**
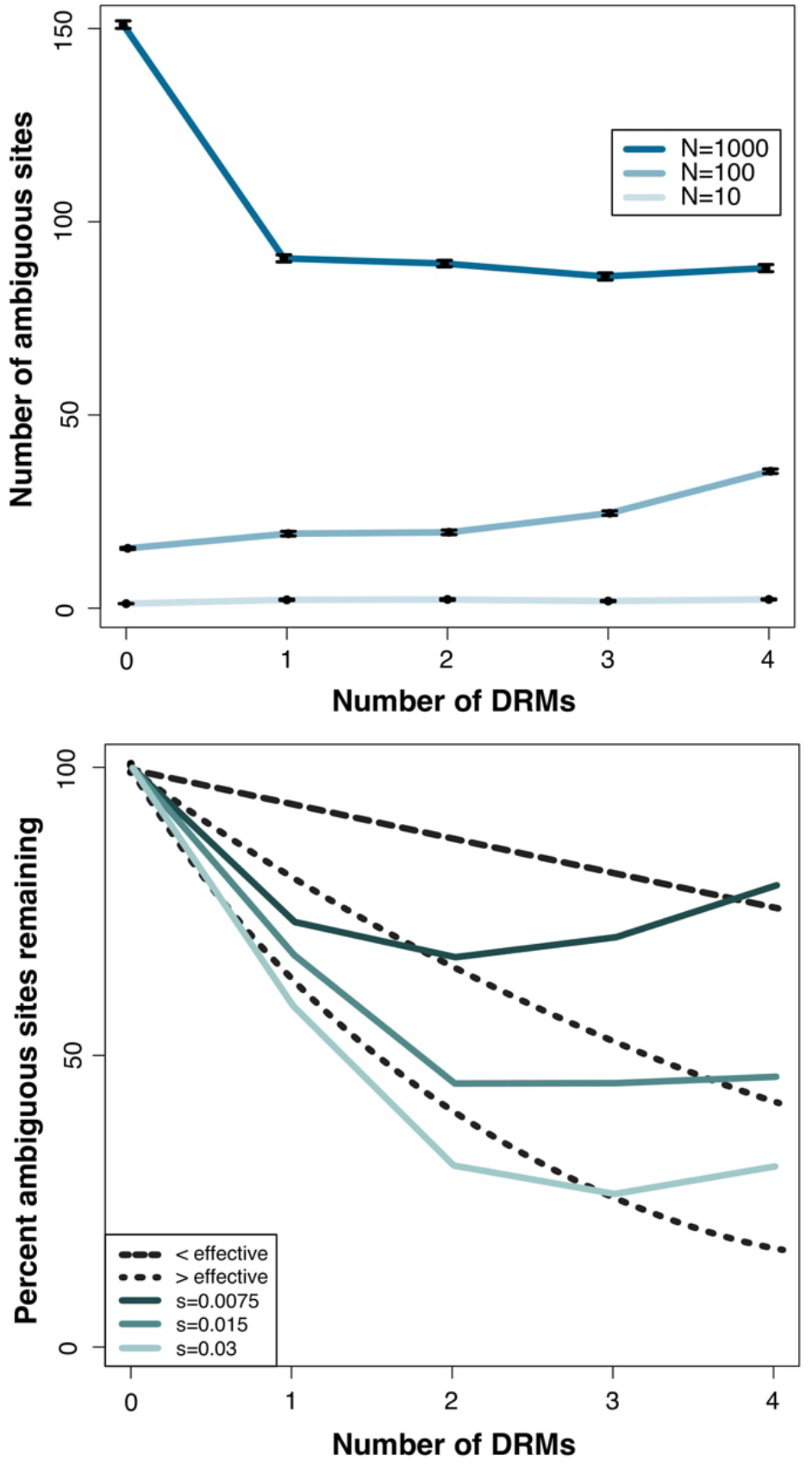
Interpreting differing levels of sequence variation in patients. Following Fig 3 of Feder *et al*. [13], the number of ambiguous sites under different models (their proxy for diversity) are plotted. In both panels, the y-axis gives the number of ambiguous sites, and the x-axis the number of drug-resistant mutations (DRMs). In panel (a) *s*=0.05 and the severity of the treatment induced bottleneck varies, with each colored line giving the census size to which the population was reduced prior to recovery. In panel (b), a non-equilibrium model is shown (a bottleneck size of 10^3^) in which the strength of selection on each DRM is varied, with the colored lines giving the corresponding selection coefficients. Overlaid in black lines are the HIV data presented in Feder *et al.* [13], with the long dashed line corresponding to two categories of less effective treatments, and the two short dashed lines corresponding to two categories of more effective treatments. As shown, both a simple bottleneck model in which the severity of size reduction is related to the efficacy of the treatment (*i.e.,* less effective treatments have less severe bottlenecks) and a hard sweep model in which the selection coefficient is related to the efficacy of treatment (*i.e*., more effective treatments are associated with larger selection coefficients), span the observed levels of variation and make it unnecessary to invoke soft selective sweeps.

Secondly, pertaining to their claim that the drug resistant phenotype is unrelated to the selection coefficient of the drug resistant genotype, one may examine whether this pattern may simply owe to a model in which the level of resistance is associated with the underlying strength of selection. To begin, given that the effect size of a selective sweep is a function of *s*/*r* (where *s* is the selection coefficient, and *r* is the recombination rate [1, 23]), it is well appreciated that selective sweeps involving mutations with large selection coefficients will reduce variation more strongly than those involving small selection coefficients. Particularly given the high recombination and mutation rates of HIV, even a small difference in the selection coefficient of a sweeping mutation may have a large effect on the number and distribution of high-frequency haplotypes that remain after the sweep is completed.

Specifically, as populations are located further from a phenotypic optimum, mutations of larger effect become possible and the variance in the effect size of fixed beneficial mutations increases (*e.g.,* [24–26]). It has been demonstrated in microbial populations that extreme environmental conditions, such as those imposed by drug treatments, can alter the underlying distribution of beneficial fitness effects (DBFE) in this manner (*e.g*., [27–29]). Multi-drug regimens impose very strong selection on HIV populations, moving wild-type populations far from the optimum. We therefore expect a high variance in the selection coefficients of the resistance mutations that fix under different drug treatments. Further, we can reasonably expect that alternative treatment strategies impose DBFEs of variable shape and scale and have different pathways to resistance, and that DRMs of more effective treatments will be on average of larger beneficial effect.

Furthermore, a substantial body of evidence indicates that non-nucleoside reverse-transcriptase inhibitor (NNRTI) resistance mutations—one of the groups of treatments categorized by Feder *et al*. as being more effective and associated with hard sweeps—may avoid pleiotropic tradeoffs in fitness, largely because the target site of NNRTIs is located far from the active site of the reverse-transcriptase enzyme [30–33]. Regardless of the differences in the selective environments of different treatments, an NNRTI resistance mutation conferring the same level of resistance as a mutation conferring resistance to another class of drug would therefore be expected to have a larger selection coefficient than resistance mutations that have tradeoffs for enzymatic function and replication rate. Additionally, NNRTI resistance requires only a single point mutation, which is expected to be associated with a larger selection coefficient compared to cases where resistance requires sequential or simultaneous fixation of several resistance mutations [31].

We compared the percent reduction in ambiguous sites (their proxy for level of variation) with zero, one, two, three, and four DRMs (the same range as analyzed by Feder *et al*. [13]) for weak and strong beneficial selection coefficients. The average percentage reductions in ambiguous calls for each additional DRM were 10.1 and 23.3, respectively, with a significantly greater reduction in diversity for the strongest selection coefficient (*p* << 0.001; Fig 3B), not surprising given that the mean sojourn time for a beneficial mutation is inversely proportional to its selection coefficient [34], giving mutations of smaller benefit a longer period of time to recombine onto additional haplotypes during a sweep. Overlaying the data of Feder *et al.* [13] onto these plots (Fig 3B), one can readily see that the expected differences in selection coefficients under hard sweep models alone (*i.e*., smaller selection coefficients under less effective treatments, larger selection coefficients under more effective treatments) re-capitulate their observation, well spanning the range in levels of diversity that they associate with hard and soft sweeps.

In sum, Feder *et al.* [13] assume identical selection coefficients and demographic histories between treatments and interpret varying levels of reduced variation as a pattern of hard and soft sweeps. However, the reasonable and well-supported expectation that the selection coefficients of drug resistant mutations will vary by drug treatment nicely accounts for the systematic differences observed, as does simply accounting for the differing bottleneck severities associated with effective vs. ineffective treatment strategies. Thus, a logical alternative appears to be that stronger treatments induce stronger bottlenecks, leading to a general reduction in variation, and impose stronger selection, leading to beneficial mutations with larger selection coefficients and a greater reduction in variation with each (hard) sweep.

## CONCLUSIONS

The above described analyses and critiques suggest a few defining features underlying recent claims of soft selective sweeps in the literature. First and foremost, classifying genomic patterns of variation in the genome of any organism into three categories under the assumption that an equilibrium neutral model will be characterized by ample variation and many rare haplotypes, that a soft sweep model will be characterized by multiple common haplotypes, and that a hard sweep model will be characterized by greatly reduced variation and a single haplotype, is inherently prone to incorrect inference when applied to real data. As has been long established, neutral population histories may well replicate these selection model expectations - a point of particular note here given that all three of the analyses investigated based their inference on genome-wide patterns of variation. For example, one can readily envision that if only given the above three model choices, a weak population bottleneck would be expected to most closely resemble pervasive soft sweeps while a strong population bottleneck would be expected to most closely resemble pervasive hard sweeps. Indeed, these expectations are met when examining the performance of these newly proposed approaches under realistic alternative models, as we have here demonstrated.

Hence, the lack of sufficient model testing and statistical performance analyses underlying these claims of recurrent soft sweeps appears to have led to inaccurate views of the evolutionary processes and trajectories governing these organisms under study. Furthermore, the generalization of these results has resulted in misleading answers to decades old questions in population genetics, with some suggesting a dominant role for positive selection in shaping patterns of genomic variation (*e.g.,* [21]). However, we suggest that studies which seek to characterize the frequency and impact of selective sweeps using population genomic data, but begin with the assumption that positive selection is the pervasive and dominant force shaping genome-wide patterns of variability, are circular to the point of being futile. In response, our work highlights the importance of first considering the demographic history of the population under study when performing genomic analyses – a history which will be non-equilibrium in any natural population. Secondly, it is crucial to evaluate Type I and Type II error pertaining to the identifiability of selection models within the context of that population history. Finally, if identifiable at all, one may consider the role of selective effects. Thus, in sum, we fundamentally argue that the characterization of positively selected genomic regions is a third step at minimum, and methodologies or analyses which rather place it as the exclusive single step are inherently prone to serious error.

## METHODS

### Implementation of the proposed methodologies

#### H-statistics

To test the robustness of the *H*-statistics (H12 and H2/H1) to demographic model misspecification, we simulated genomic data using the backward-in-time simulator *msms [35]*. To match the simulations of Garud *et al.* [10], we sampled 145 individuals belonging to a population sample simulated under a neutral mutation rate of 1x10^-9^ events/bp/gen, and a recombination rate of 5x10^-7^ cM/bp. Simulated loci were 10^4^ bp in length, concordant with Garud *et al*.’s Fig S11, which shows that the 400 SNP windows used to calculate empirical DGRP *H*-statistics were, on average, consistent with 10^4^ bp genome segments. Furthermore, using the aforementioned mutation and recombination rates, combined with the demographic parameters described below, the mean number of segregating sites (~16,000 replicates) was 406 for neutral simulations and ~380 for sweep simulations.

Using these parameters, we simulated neutrality, soft sweeps, and hard sweeps. For sweep simulations, the positively selected site (selection coefficient ~*U*[0, 1]) was positioned in the middle of each locus, as this is an assumption of both *H*-statistics (albeit an unreasonable and best-case scenario, inasmuch as it assumes that the target of selection is known *a priori*). Similar to Garud *et al*. [10], we considered sweeps generated from a beneficial mutation rate, where hard sweeps had a low recurrent beneficial mutation rate of *Smu* = 0.01 and soft sweeps had a higher rate of *Smu* = 10. In all cases, and in order to be consistent between models, we conditioned on the start time of a sweep.

To fully explore the 95% credible interval of the DGRP admixture model, we generated 16,000 neutral simulations drawing parameters randomly from the posteriors given in Duchen *et al.*’s Fig S10 (and pers. comm., reported here in S1 Table) [15]. For comparison to Garud *et al*’s [10] findings, we also generated hard and soft sweep simulations (16,000 each) from the posterior parameter values and sampled sweep starting times from the time of admixture until the present. All *msms* commands are documented in S1 Text.

To quantify the fit of each of the top 50 empirical H12 peaks to a simulated distribution, we calculated empirical *p*-values. The neutral H12 *p*-value represents the proportion of neutral simulations that are more extreme than the empirical H12 values. Garud *et al*.’s test is designed to be a two-step approach, where first H12 outliers are identified and then H2/H1 values are used to determine the type of sweep. Therefore, we determined the range of the 2.5% largest neutral H12 values. We then removed all simulations with H12 values smaller than the lower neutral bound to calculate empirical H2/H1 *p*-values.

#### S/HIC

S/HIC classifies genomic regions into discrete categories (hard, hard-linked, soft, soft-linked, or neutral) by comparing test (empirical or simulated) data to simulated training data. Because demographic perturbations may mask the genomic signals of selection, a user must first choose an appropriate demographic model under which to simulate training data. Using this training set, the S/HIC pipeline then estimates a range of population genetic summary statistics and implements a type of supervised machine learning called the Extremely Randomized Trees (ERT) classifier [36]. Like the more well-known Random Forests (RF), ERT aggregates the results of an ensemble of simple estimators, or decision trees. Whereas single decision trees have a tendency to over-fit to the training data, ensemble methods work by combining multiple weak classifiers to build a stronger one. For these methods to work, a forest of random, unique trees needs to be generated. RF and ERT generate randomness in different ways: while RF modifies the training set for the construction of each tree (bagging), ERT instead introduces randomness into the node splitting process and uses the entire training data set [37]. For our purposes, we made no attempt to modify or critique the S/HIC pipeline *per se*. Indeed, recent studies have shown ERT methods to generally perform as well as other RF methods [38]. Instead, our goal was to explore the robustness of S/HIC to demographic and selection model mis-specification (as is the case in all natural population analysis), and explore a wider range of demographic models, in order to understand the behavior of the statistic.

Unless stated otherwise, we replicated the simulations used in the original methods paper (see [12], S1 Table) and the proceeding paper focused upon the analysis of human data (see [11]). Briefly, training and testing sets were compiled by simulating chromosomes in *discoal* [39]. To emulate the patterns seen in regions linked to selective sweeps, thousands of replicate simulations were conducted where each chromosome was divided into 11 sub-windows with the beneficial mutation occurring in the center of a single sub-window (see [12], Fig 1). A focal hard or soft sweep was defined as occurring in the central (6^th^) window. From these simulated datasets, we used the S/HIC pipeline to calculate summary statistics, train classifiers, and classify test simulations.

Of critical importance is our correct implementation of the S/HIC method. To assess this, we first generated two equilibrium training and testing sets identical to that used to generate Schrider and Kern’s Fig 4 [12] and compared the resulting classification probabilities. In two independent trials, we correctly classified neutrality 74% and 81% of the time (compared to their 82%), and soft sweeps 80% and 75% of the time (compared to their 79%). It should be noted that this considerable variance in classification probabilities is inherent to the underlying randomness of this machine learning approach.

Next, we evaluated the robustness of S/HIC to demographic mis-inference. To understand the effects of changing the DFE in humans, we chose to reanalyze the demographic model of the East African LWK population because of its high true positive rate (see Schrider and Kern Fig.S1), likely owing to a history of comparatively small population fluctuations. We first constructed a model from the same feature vectors used to train Schrider and Kern’s LWK model. We then simulated a test set and compared our classification probabilities to ensure an ability to replicated their results. Finally, we generated another test set and lowered the strength of selection (original: 2*Ns* ~ *U*[166, 3333] (SK Table S5); new: 2*Ns* ~ *U*[10, 1000]).

We further evaluated Schrider and Kern’s less severe bottleneck model (population size reduced to 29% at 0.044 4*N* generations, recovering at 0.0084 4*N* generations) as the true model, and quantified the ability to correctly classify neutral loci simulated under more severe bottlenecks. To address models of population structure, we simulated data sets with two populations diverging at 0.25, 0.5, or 1.0 in 4*N* generations, followed by isolation. We then sampled 90 individuals from one population and 10 from the other, in order to investigate the effects of undetected population structure. Finally, we considered migration models consisting of pulses (5, 10, 50, or 100) of migration (0.00001, 0.0001, 0.001, 0.01, or 0.1% of the population) occurring every 10 generations until the present, following the initial split. These datasets were then classified using the equilibrium dataset.

All *discoal* commands are given in S1 Text.

### Simulation of HIV populations

Simulated populations of HIV were generated via forward-in-time population genetic simulation in the *SLiM* version 3 software package [40]. We simulated sequences of 1980 nt (equivalent to the combined lengths of the reverse-transcriptase and protease genes) with demography and positive selection approximating the stages of 1) initial infection, 2) population growth, 3) a treatment-induced population bottleneck, and 4) sequential fixation of drug resistant mutations (DRMs). Specifically, populations were initiated from a single infecting virion [41,42] and then grew exponentially for 100 generations until reaching a population size of 10^4^. After 1,000 generations of neutral evolution, a population bottleneck to either 10^3^, 10^2^, or 10 individuals was imposed. The population mutation rate was set to 3.4x10^-5^ per site per replication [43], and the recombination rate was 1.4x10^-5^ per site per replication [44].

Beneficial mutations of effect size *s* = 0.03, *s* = 0.015, and *s* = 0.0075 (corresponding to weak, moderate, and strong selection) fixed sequentially until a specified number of DRMs were present in the population (zero, one, two, three, or four) with the population size recovering in a stepwise manner as each DRM fixed to reflect sequential fitness gains. When the desired number of DRMs was achieved, the population size was fully restored to its pre-treatment value of 10^4^ (*i.e*., the population size recovered once resistance was achieved) and the population was allowed to evolve neutrally for 100 generations before analysis. Mutations were introduced to the population in the following manner: every one or five generation/s (for the weak and strong selection regimes, respectively), a new copy of a beneficial mutation was introduced if no beneficial mutation was presently segregating in the population. For the purposes of the simulations, beneficial mutations were considered to be fixed once they reached a frequency of 0.95 or greater. Our populations therefore experienced selective sweep dynamics, wherein each DRM only fixed in the population after the previous sweep was at or near completion. The rarity of new beneficial mutations was scaled with the strength of selection and population bottleneck size, with the assumption being that the rate of beneficial mutations is a product of the population size and the beneficial mutation rate appropriate for a given specific effect size.

At the end of the recovery period following sequential DRM fixation, and following Feder *et al*. [13], the number of ambiguous base calls for the population was approximated as the number of sites with a minor allele frequency greater than or equal to 0.15, assuming this as a reasonable threshold at which population-level Sanger sequencing would result in an ambiguous base call at the site. The number of ambiguous base calls was estimated for populations experiencing different strengths of selection and variable strengths of treatment-induced bottleneck.

Finally it is noteworthy that variation in the ∆DRM measurement of Feder *et al*. [13] within treatment categories does not always correlate strongly with their measurement of treatment effectiveness. In fact, the unboosted PI treatments of the greatest and second-least effective treatment have the least and greatest DRM-associated decreases in diversity, respectively, counter to their central argument.

## ACKNOWLEDGEMENTS

We thank Claudia Bank, Brian Charlesworth, Matt Jones, Stefan Laurent, Mike Lynch, Susanne Pfeifer, Wolfgang Stephan, Lucy Tran, Miltos Tsiantis, Alex Wong, and three anonymous reviewers for helpful comments on the manuscript. We further thank Dan Schrider for assistance with the implementation of S/HIC, and Pablo Duchen and Stefan Laurent for assistance with the Drosophila *msms* simulations and for providing posterior distributions under their model.

## Supporting Information

**S1 Fig.**
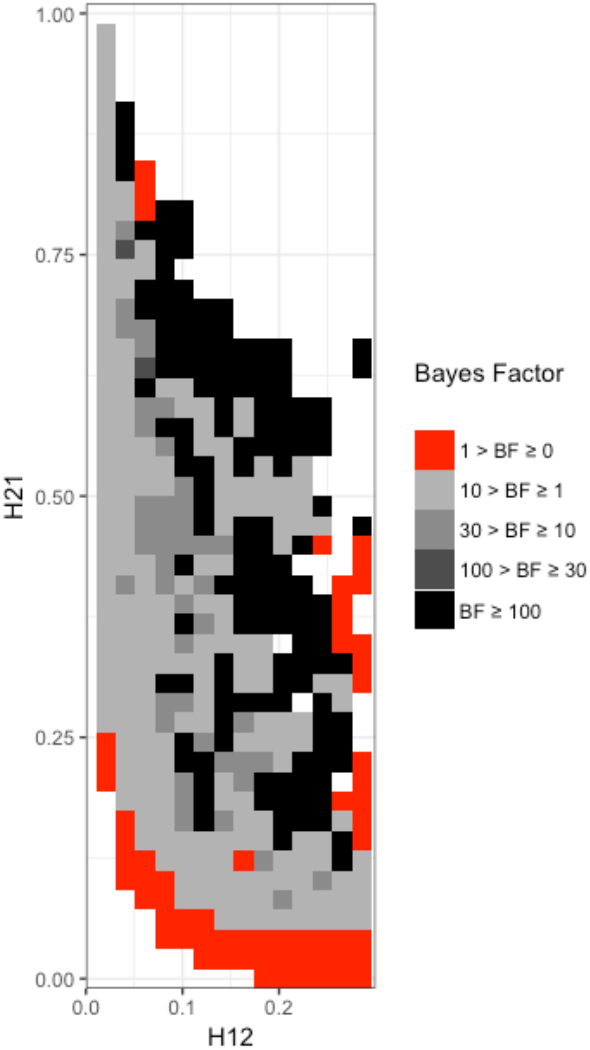
Range of H12 and H2/H1 values expected under hard and soft sweeps. We depict our results in the same fashion as Garud *et al*. (their Fig 11) to demonstrate how plotting of Bayes factors (BFs) can lead to erroneous conclusions about the ability of the *H*-statistics to differentiate between hard and soft sweeps. BFs were calculated by taking the ratio of the number of soft sweep versus hard sweep simulations that were within a Euclidean distance of 10% of a given pair of H12 and H2/H1 values. Red portions of the grid represent H12 and H2/H1 values that are more easily generated by hard sweeps (generated with *Smu* = 0.01), while grey portions represent regions of space more easily generated under soft sweeps (generated with *Smu_A_* = 10). These results are based on 16,000 hard and soft sweep simulations.

**S2 Fig.**
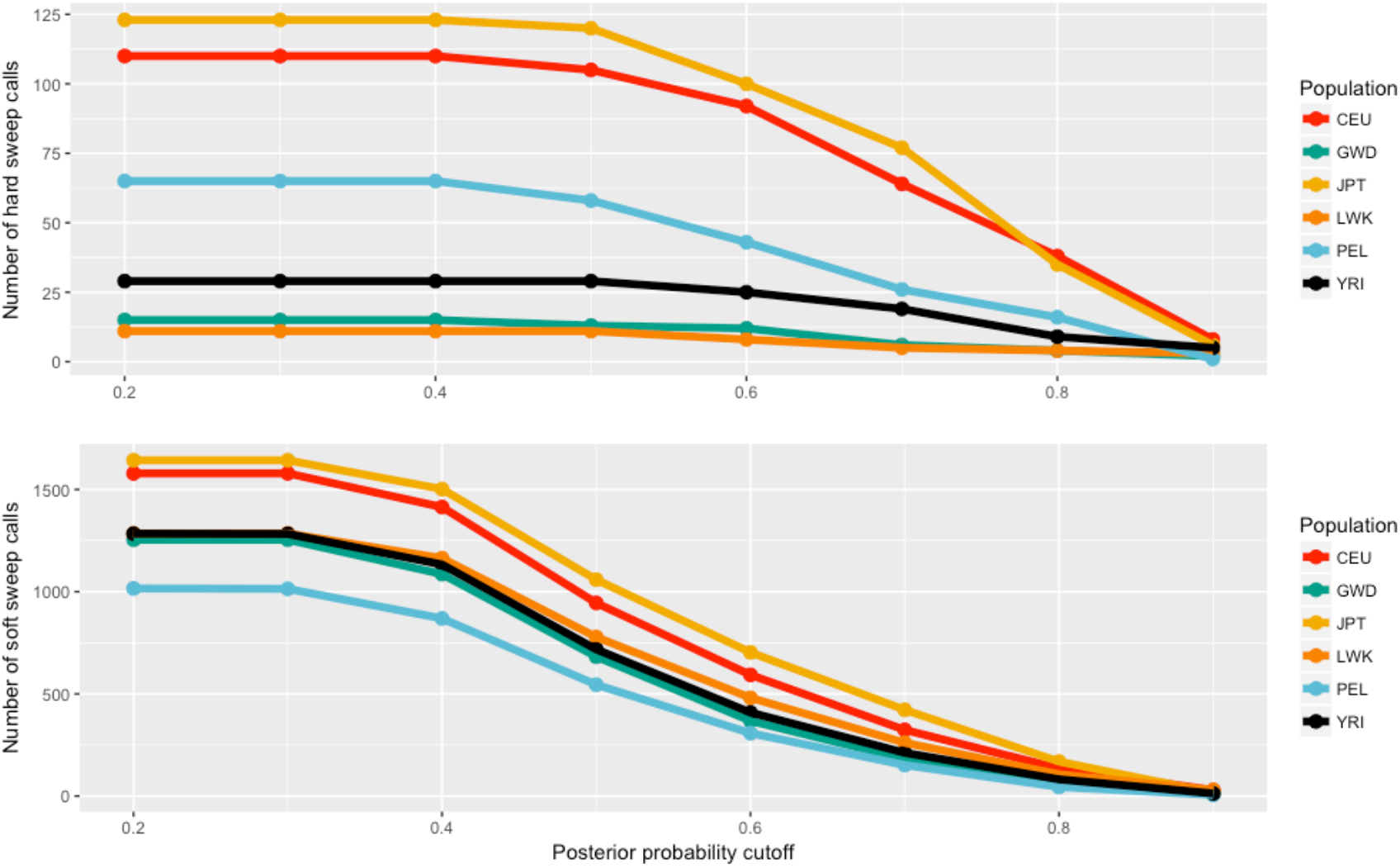
Impact of using a posterior probability threshold in the classification of sweeps using S/HIC. Drop-off in the number of genomic regions classified as hard (top) and soft (bottom) sweeps as a posterior probability threshold is imposed. Data plotted from Schrider and Kern’s S2 Table.

**S3 Fig.**
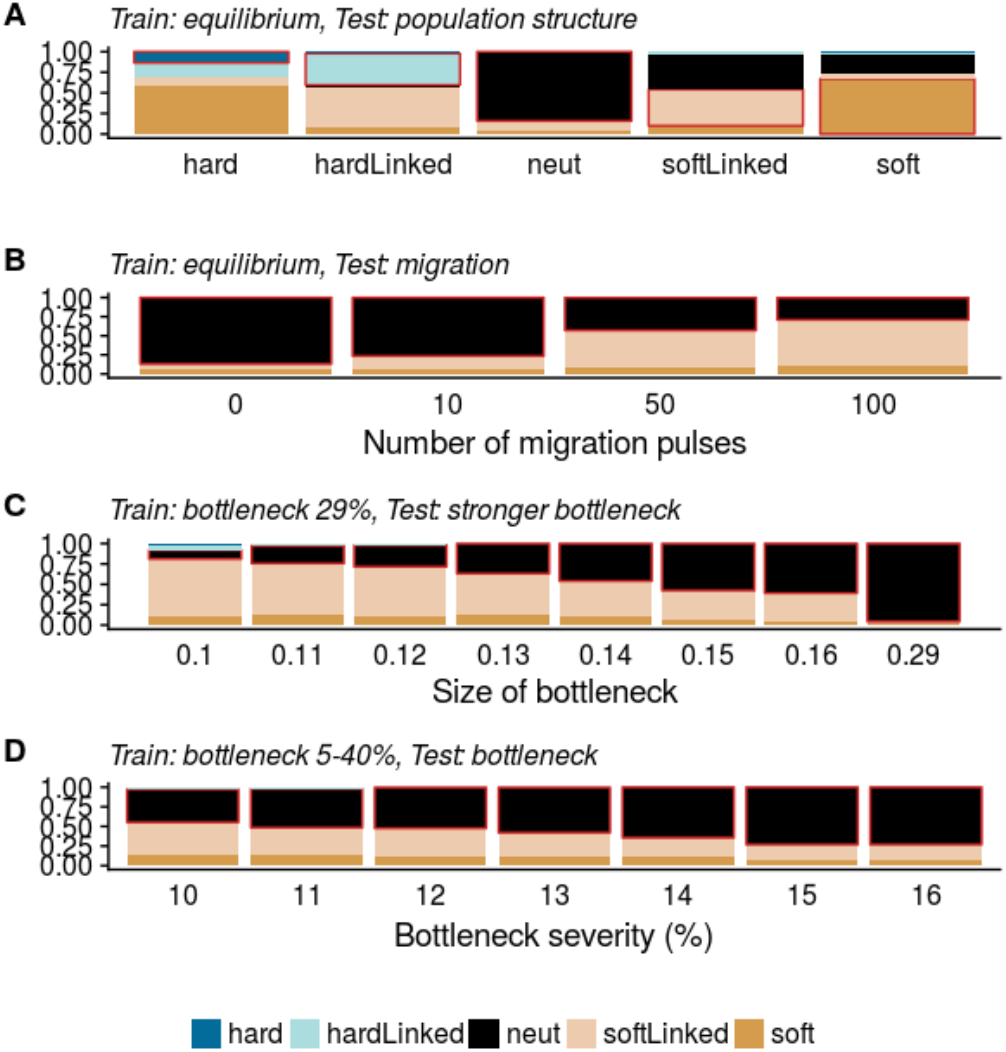
S/HIC classification performance under mis-specified demography. (a) Both test and training data were simulated under a constant size demographic model; however, the test data set consists of 100 individuals sampled from two populations (9:1 sampling ratio) which diverged 0.50 x 2*N* generations in the past. (b) The classification performance when neutrality is simulated under a structured, constant size population with migration, varying the number of pulses of gene flow. Here, the populations diverged 4*N* generations ago, with 10 migrants per pulse. (c) The classification performance of neutrality simulated under a bottleneck model with varying bottleneck severity (*e.g*., a ’size of bottleneck’ of 0.1 corresponds to a temporary reduction to 10% of the ancestral size), when the true model is a bottleneck decreasing the population to 29% of the ancestral size. Bottleneck timing and duration are consistent across all test and training sets, with the bottleneck beginning at 0.044*4*N* generations ago and returning to the initial population size 0.0084*4*N* generations ago. (d) Similar to (c), however the training set was constructed from a range of bottleneck simulations which decrease the population to 5-40% of the ancestral size.

**S1 Table. Posterior distribution of parameter estimates from the DGRP admixture model of Duchen *et al*.**

**S2 Table. Neutral migration models investigated, and the resulting S/HIC classifications.**

**S1 Text. Simulation code.** *msms* commands used to simulate datasets for calculating *H*-statistics and *discoal* commands used to simulate data sets for S/HIC.

